# FDX1 regulates cellular protein lipoylation through direct binding to LIAS

**DOI:** 10.1101/2023.02.03.526472

**Authors:** Margaret B. Dreishpoon, Nolan R. Bick, Boryana Petrova, Douglas M. Warui, Alison Cameron, Squire J. Booker, Naama Kanarek, Todd R. Golub, Peter Tsvetkov

## Abstract

Ferredoxins are a family of iron-sulfur (Fe-S) cluster proteins that serve as essential electron donors in numerous cellular processes that are conserved through evolution. The promiscuous nature of ferredoxins as electron donors enables them to participate in many metabolic processes including steroid, heme, vitamin D and Fe-S cluster biosynthesis in different organisms. However, the unique natural function(s) of each of the two human ferredoxins (FDX1 and FDX2) are still poorly characterized. We recently reported that FDX1 is both a crucial regulator of copper ionophore induced cell death and serves as an upstream regulator of cellular protein lipoylation, a mitochondrial lipid-based post translational modification naturally occurring on four mitochondrial enzymes that are crucial for TCA cycle function. Here we show that FDX1 regulates protein lipoylation by directly binding to the lipoyl synthase (LIAS) enzyme and not through indirect regulation of cellular Fe-S cluster biosynthesis. Metabolite profiling revealed that the predominant cellular metabolic outcome of FDX1 loss-of-function is manifested through the regulation of the four lipoylation-dependent enzymes ultimately resulting in loss of cellular respiration and sensitivity to mild glucose starvation. Transcriptional profiling of cells growing in either normal or low glucose conditions established that FDX1 loss-of-function results in the induction of both compensatory metabolism related genes and the integrated stress response, consistent with our findings that FDX1 loss-of-functions is conditionally lethal. Together, our findings establish that FDX1 directly engages with LIAS, promoting cellular protein lipoylation, a process essential in maintaining cell viability under low glucose conditions.

## Introduction

Ferredoxins (FDXs) are small iron-sulfur cluster containing proteins that mediate electron shuttling among multiple metabolic pathways. FDXs generally act as electron donors and are essential in a wide array of fundamental reactions including iron–sulfur cluster and steroid hormone biosynthesis and other cytochrome-P450-associated reactions (1). Humans have two ferredoxins, FDX1 and FDX2. In human cells FDX1 was shown to regulate steroid hormone synthesis by donating electrons to various cytochrome P450 proteins such as CYP11A1 (2–4). This function is executed only in steroidogenic tissues explaining the particularly high mRNA levels of FDX1 in human adrenal glands (5,6). FDX2 on the other hand was shown to be a central component of cellular iron-sulfur cluster biosynthesis (7–11). Much of the early studies on human ferredoxin function were conducted using bacteria or yeast reconstitution models or in vitro functional assays. As these experimental approaches did not necessarily explore the natural function of FDX1 within the human cell it produced incomplete and sometimes contradicting findings on the contribution of human ferredoxins and FDX1 in particular to Fe-S cluster (9,10,12,13) and Heme biosynthesis (14,15) and more recently in the regulation of protein lipoylation (13,16). As such, the predominant natural role(s) of FDX1 in non-steroidogenic human cells remains poorly characterized.

We recently reported FDX1 as a crucial regulator of copper dependent cell death (cuproptosis) (16,17). Copper ionophores, such as elesclomol, bind and shuttle Cu (II) to the mitochondria where the Cu (II) can be reduced to Cu(I) by FDX1 promoting the cytotoxic effect of copper (17). Copper then induces the destabilization of Fe-S cluster proteins and can directly bind lipoylated proteins promoting their aggregation and ultimately leading to cell death (16). Interestingly, we revealed that FDX1 loss-of-function in cancer cells also results in lack of protein lipoylation and that FDX1 protein levels are highly correlated with overall levels of lipoylated protein levels across hundreds of different cancer types (16). Thus, establishing that FDX1 is an upstream regulator of protein lipoylation in human cells, a finding that was recently corroborated (13).

Protein lipoylation is a lipid post-translational modification that occurs on conserved lysine residues of only four key mitochondrial metabolic complexes. Lipoylation is essential for the enzymatic function of the lipoylated complexes which include the pyruvate dehydrogenase (PDH), α-ketoglutarate dehydrogenase (KGDH), branched-chain α-ketoacid dehydrogenase (BCKDH), and glycine decarboxylase complex (GDC), also referred to as the glycine cleavage system. This lipoylation process is evolutionarily conserved from bacteria to humans and has been mostly characterized in bacteria and yeast systems and less so in human cells. In humans, lipoic acid is synthesized de novo in a multi-step process. First, an 8-carbon fatty acid (octanoic acid) synthesized in the mitochondria covalently attached to an acyl-carrier protein (ACP), is transferred to a conserved lysine residue within the Glycine Cleavage System Protein H (GCSH) protein by lipoyl (octanoyl) transferase 2 (LIPT2). Second, two sulfurs are added at C-6 and C-8 positions by the lipoyl synthase (LIAS) enzyme to form the lipoylated form of GCSH. In this reaction the sulfur atoms are provided by the auxiliary Fe-S cluster of LIAS, which requires Fe-S cluster reconstitution on LIAS after each catalytic cycle of lipoic acid synthesis (18–20). Therefore, any perturbation in cellular Fe-S cluster biosynthesis will result in reduced ability of the cell to replenish the auxiliary Fe-S cluster of LIAS after each cycle of lipoic acid synthesis, and ultimately reduced levels of protein lipoylation. Given the contradicting findings on the role of FDX1 in Fe-S cluster biosynthesis and our observation that FDX1 regulates protein lipoylation (16) we set out to determine the predominant role of FDX1 in human cells and characterize the transcriptional and metabolic outcome following FDX1 loss-of-function.

## Results

### FDX1 regulation of lipoylation is not mediated by Fe-S cluster availability

In steroidogenic tissues FDX1 plays a unique role in the process of steroid hormone biosynthesis by contributing electrons to P450 complexes such as CYP11A1 (6). However, while most cells previously characterized for gene expression as part of the Cancer Cell Line Encyclopedia (CCLE) show no detectable CYP11A1 mRNA levels (Fig. S1A), FDX1 is uniformly expressed across all cell lines from different lineages (Fig. 1A). This suggests that in most human cells FDX1 has a function beyond steroid hormone biosynthesis. To better understand the function of FDX1 in these cancer cells we used the Cancer Dependency Map (www.depmap.org) resource that measures the viability effect of genome-wide CRISPR/Cas9 loss-of-function in each individual cell line across more than a thousand different cell lines. Using this resource, gene function can be inferred by correlating the effect on viability of a specific gene loss-of-function (dependency) to the dependencies of all other genes across all examined cell lines. Gene dependencies with similar viability profiles across hundreds of cancer cell lines are expected to fall within the same pathway. This approach has been successfully applied by numerous groups to reveal unknown functions of genes of interest (21–24). Using this approach, the top correlate of FDX1 dependency is LIAS, the only gene known to synthesize the lipoyl moiety in the mitochondria (Fig. 1B). This correlation between the essentiality of FDX1 and LIAS is highly significant (Fig. 1C) and the reciprocal analysis shows that FDX1 is also within the top correlates of LIAS loss-of-function (Fig. S1B). Other enzymes in the lipoylation pathway (GCSH, LIPT1, DLD, LIPT2) also scored high and analysis of enriched functional categories in the top 1,000 genes correlating with FDX1 dependency revealed that protein lipoylation is the most enriched category (Fig. 1D, table S1). These results were unique to FDX1, as a similar analysis for its paralog gene FDX2 did not result in any gene in the lipoylation pathway scoring in the top 1,000 co-dependent genes (Fig. 1E, table S1). Moreover, the top correlates for FDX2 were genes in the Fe-S cluster biosynthesis pathways such as HSCB and LYRM4 and the ferredoxin reductase (FDXR) enzyme. This genetic-perturbation data analysis approach first establishes FDX1 loss-of-function is a selective dependency with most cells showing a slight growth impairment. Second, it shows that FDX1 and FDX2 differ in their function; the predominant predicted role of FDX1 is associated with cellular protein lipoylation pathway, whereas the role of FDX2 is predicted to be involved in Fe-S cluster biosynthesis.

**Fig. 1.**
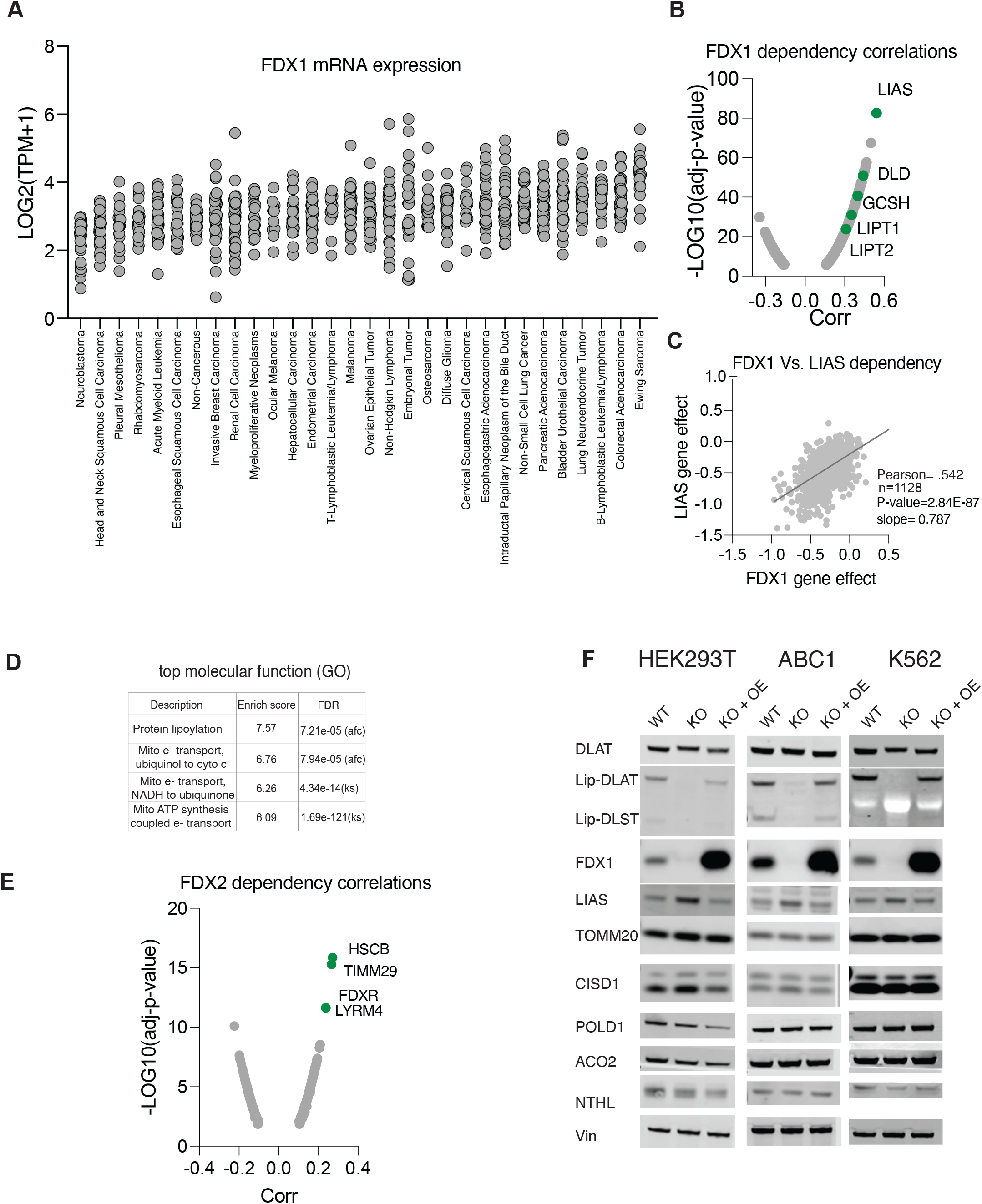
Genetic evidence supporting the role of FDX1 in regulating protein lipoylation. (A) FDX1 mRNA expression levels in cancer cell lines from the CCLE categorized by lineage. (B) The CRISPR/Cas9 FDX1 loss-of-function viability outcome in 1128 cell lines correlated to the loss-of-function of all other tested genes. Plotted are the correlation values and the p-value for the 1000 most significant correlations. In green are genes associated with the protein lipoylation pathway. (C) The correlation of LIAS and FDX1 CRISPR/Cas9 loss-of-function across 1128 cell lines. (D) Top molecular function (GO) enriched categories in the 1000 genes that have their loss-of-function most correlated with FDX1 loss-of-function. (E) Indicated proteins were analyzed using immunoblot assays of HEK293T, K562 and ABC1 WT cells or cells with CRISPR/Cas9 induced FDX1 loss-of-function (KO) alone or with reconstituted FDX1 (KO+OE). The data used to generate plots A-D is provided as supplementary table 1.

To experimentally characterize the role of FDX1 we used CRISPR/Cas9 in ABC1, HEK293T and K562 cells to induce FDX1 loss-of-function (FDX1 KO) and controlled for off-target effects by overexpressing FDX1 in the FDX1 loss-of-function cells (KO+OE). We did this in three different cell lines (K562, ABC1, and HEK293T) to establish the FDX1 dependent phenotypes. In all three cell lines protein lipoylation was completely abolished upon FDX1 loss-of-function (KO) and completely restored upon FDX1 reconstitution (KO+OE) (Fig. 1F). This is not due to changes in total protein levels of the lipoylated proteins (as shown for DLAT). Cellular protein lipoylation is regulated by the Fe-S cluster enzyme LIAS (25,26) that uses the sulfurs from its auxiliary Fe-S cluster to facilitate the double sulfur insertion to the sixth and eighth carbons of the octanoyl precursor of lipoic acid (18). This dependency of LIAS on Fe-S clusters for function has been shown in genetic models where perturbations to Fe-S cluster biosynthesis results in reduced levels of LIAS and consequently reduced levels of protein lipoylation (27,28). However, in the FDX1 KO cells LIAS levels were not reduced, and stabilization of LIAS was observed in FDX1 KO cells as compared to WT and OE cells (Fig. 1F). This suggests that FDX1 does not regulate LIAS activity by reducing Fe-S cluster availability but rather by an upstream process that inhibits the function of LIAS, preserving its auxiliary Fe-S cluster and resulting in its stabilization. We further confirmed this model by showing that FDX1 protein levels had no effect on the stability of other Fe-S cluster containing proteins such as CISD1, POLD1, ACO2, and NTHL (Fig. 1F).

### FDX1 regulates cell respiration and ability to proliferate in low glucose conditions

Lipoylation regulates the activity of key carbon entry points into the TCA cycle by directly regulating the activity of α-ketoacid dehydrogenases, a family of multi-subunit enzyme complexes that includes PDHC, OGDC, and branched-chain ketoacid dehydrogenase complex (BCKDC) (26). Consistent with the observed effect on protein lipoylation, FDX1 KO cells exhibit low oxygen consumption that is completely restored upon FDX1 reconstitution (Fig. 2A-C). Despite the profound effect on cellular respiration only mild growth impairment in FDX1 KO cells is observed when cells were cultured in 10mM glucose conditions (Fig. 2D). However, when glucose levels were reduced to 5mM (normal physiological levels) or 2mM (low physiological level), FDX1 KO cells exhibited a strong growth defect consistent with FDX1 being essential for mitochondrial metabolism (Fig. 2D and S2). Together, our data show that FDX1 is an essential upstream regulator of protein lipoylation and shows metabolic conditional lethality, essential in cell states where cells rely on mitochondrial energy metabolism to sustain proliferation.

**Fig. 2.**
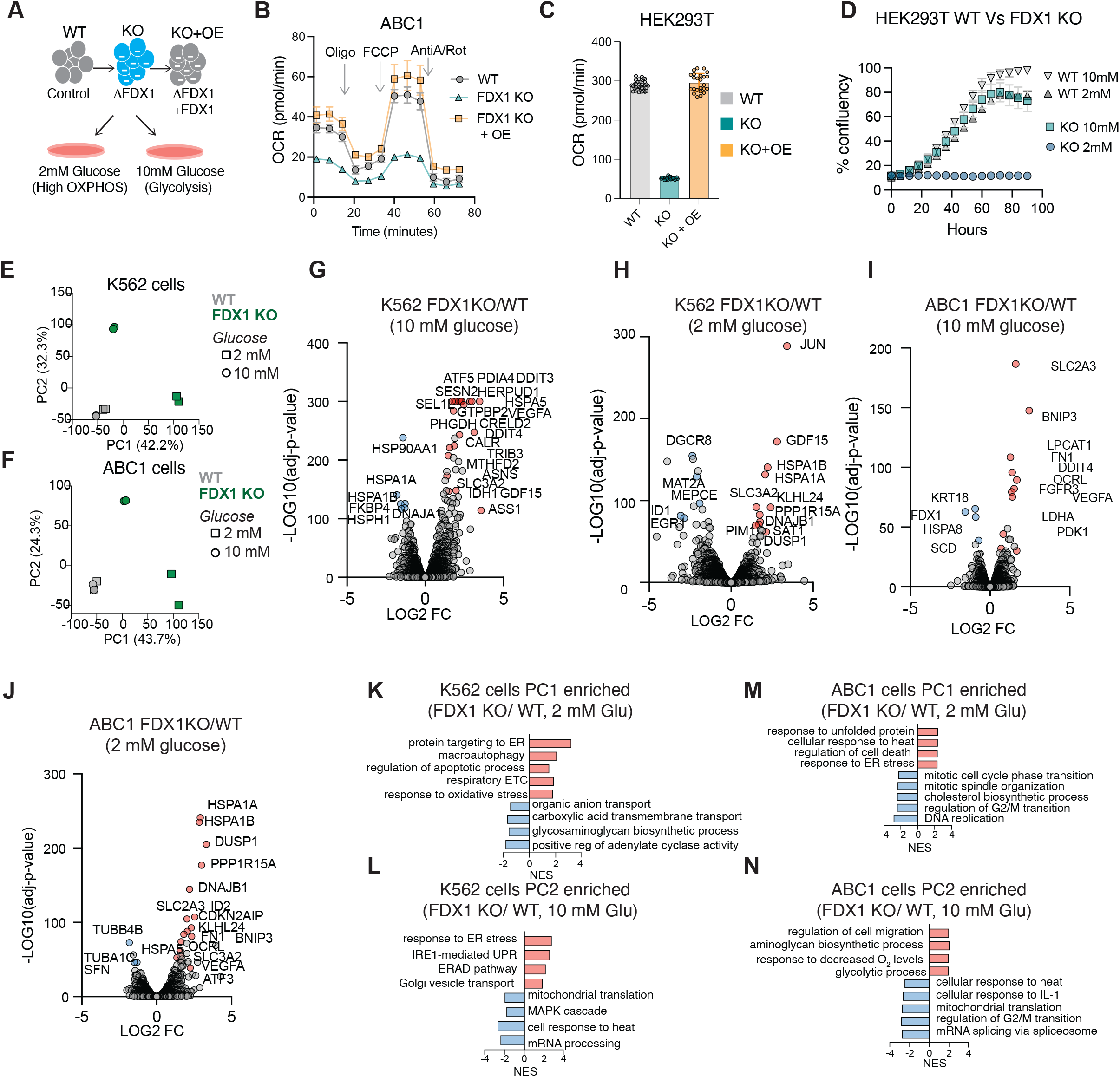
FDX1 loss-of-function induces the integrated stress response and inhibits proliferation and respiration in low glucose conditions. (A) schematic of the experimental conditions. (B) The oxygen consumption rate (OCR) was detected in ABC1 WT, FDX1 KO and FDX1 KO cells with FDX1 expression reconstituted and after the addition of oligomycin (ATP-linked), the uncoupler FCCP (maximal), or the electron transport inhibitor antimycin A/rotenone (baseline) (mean ± SD, of at least 4 biological replicates). (C) Basal OCR was measured in HEK293T WT, FDX1 KO and FDX1 KO cells with FDX1 expression reconstituted (mean ± SD, of at least 8 biological replicates). (D) Cell proliferation as measured by confluency is shown for HEK293T WT and FDX1 KO cells grown in either 10mM or 2mM glucose containing media. (n=3) (E-F) PCA of gene expression levels derived from RNA-seq in ABC1 (E) and K562 (F) WT (gray) and FDX1 KO (green) cells growing in either 2mM (square) or 10mM (circle) glucose conditions (n=2 for each condition). (G-J) Volcano plots relating to LOG2 fold-change (x-axis) and - LOG10(p-values) (y-axis) for gene expression changes observed in FDX1 KO as compared to WT. Volcano plots for K562 (G, H) and ABC1 (I, J) show cells growing in media containing either 10mM (G, I) or 2mM (H,J) glucose. (K-N) Gene set enrichment analysis (GSEA) of GO_Biological Processes using either PC1 (K, M) or PC2 (L, N) weights as the gene ranks. Representative categories were selected, and full data is provided in Table S2.

### The transcriptional changes associated with FDX1 loss-of function

To enable better characterization of the FDX1 KO cell state we performed global gene expression analysis using a bulk RNA-seq approach. We profiled both WT and FDX1 KO cells growing in either 10mM glucose (where FDX1 is non-essential) and 2 mM glucose (where FDX1 is essential) conditions. This analysis was done in both ABC1 and K562 cells. PCA of the gene expression in the different conditions revealed a similar trend between the different cell lines. The first principal component (PC1; explaining 42.2% and 43.7% of variance in K562 and ABC1 cells respectively) was largely driven by the deleterious effect of FDX1 loss-of-function under low glucose conditions (2mM). The second principal component (PC2; explaining 32.3% and 24.3% of variance in K562 and ABC1 cells respectively) was associated with FDX1 loss-of-function in normal growth conditions where the perturbation results in inhibition of proliferation rate but is not essential (Fig. 2E-F). In both cell lines we see a similar gene signature induction that is promoted by FDX1 loss-of-function. This includes the induction of both metabolism related genes and integrated stress response (ISR) genes (Fig. 2G-N and Table S2).

When both K562 and ABC1 cells were grown in normal 10mM glucose conditions, FDX1 loss-of-function resulted in dysregulation of metabolic genes. In ABC1 cells, there is an induction of the PDK1 (Pyruvate Dehydrogenase Kinase 1) gene alongside LDHA (lactate dehydrogenase) suggesting a suppression of the dysfunctional PDH complex and funneling of the pyruvate to the production of lactate by LDHA (Fig. 2I). There is also dysregulation of lipid metabolism genes such as SCD (downregulation) and LPCAT1 (upregulation). The induction of LPCAT1 (lysophosphatidylcholine acyltransferase 1), an enzyme that can convert lysophosphatidylcholine (LPC) to phosphatidylcholine (PC), suggests lipid homeostasis dysregulation related to PC. This is further supported by the metabolomics data below that show impairment in the de novo PC synthesis by the Kennedy pathway (Fig. 2I). In K562 cells there is a profound increase in genes related to serine and glycine metabolism such as PHGDH and MTHFD2 and TCA cycle related genes including ASS1 (argininosuccinate synthase 1) and IDH1 (isocitrate dehydrogenase 1) that all might be induced to compensate for the inactivated lipoylated complexes (Fig. 2G). When cells are shifted to low glucose conditions (2 mM), where FDX1 loss-of-function is essential, an FDX1-dependent cell stress signature is observed that includes both heat-shock genes (HSPA1A and HSPA1B) and ISR genes such as the metabokine GDF15 that is responsive to mitochondrial dysfunction (29,30) (Fig. 2H and 2J).

To better understand the specific gene categories that are most up- and down-regulated in each tested condition, we performed gene set enrichment analysis (GSEA) (31) of GO Biological processes (32) using both the PC1 and PC2 (separately) weights as the gene ranks in each cell line (see Materials and methods for details). This approach revealed that FDX1 loss-of-function resulted in a strong unfolded protein response, ER-stress gene signature consistent with the induction of the integrated stress response as previously described in the case of mitochondrial dysregulation (33). Whereas the ER-stress related genes are induced in K562 cells even when cells are grown in normal media conditions; in ABC1 cells this gene signature was observed only in low glucose conditions (Fig. 2K-N). This suggests that tolerance to FDX1 loss-of-function between different cell lines, as detected by the induction of the ISR or cell proliferation inhibition, differ between cell lines, and depend on the natural metabolic wiring of each cell type and environmental metabolite availability.

### The effect of FDX1 loss-of-function on cellular metabolism

As we show above, FDX1 is essential for cells that rely on mitochondrial metabolism. To establish that this is due to its role in regulating LIAS-mediated protein lipoylation we profiled the metabolic consequences of FDX1 loss-of-function in three different cell lines cultured in both normal and low glucose conditions. We hypothesized that the major metabolic changes would result from dysfunctional lipoylated complexes (Fig. 3A), and that these will reflect the aberrant mitochondrial metabolism in FDX1-depleted cells providing the etiology for the conditional essentiality of this gene in low glucose. We tested our hypothesis by conducting both intracellular and media metabolite profiling of WT, FDX1 KO and FDX1 KO with reconstituted FDX1 (KO+OE) cells. These experiments were conducted in ABC1, HEK293T and K562 cell lines to ensure the observed effects are not lineage specific (Table S3). Indeed, high levels of pyruvate (Fig 3B and C) and serine (Fig. 3B and D) in the media of FDX1 KO cells corroborated malfunction of PDH and GCS respectively. Pyruvate is the direct substrate of PDH, and therefore is expected to accumulate when PDH is not functional. Impaired glycine cleavage system (GCS) is expected to result in accumulation of serine converted from glycine by the mitochondrial SHMT2 enzyme (34). Moreover, dysfunctional GCS will inhibit the synthesis of 10-formyl THF which is the formyl donor for formyl-methionine (35,36) explaining the observed increased uptake of FDX1 KO cells of formyl-methionine from the media.

**Fig. 3.**
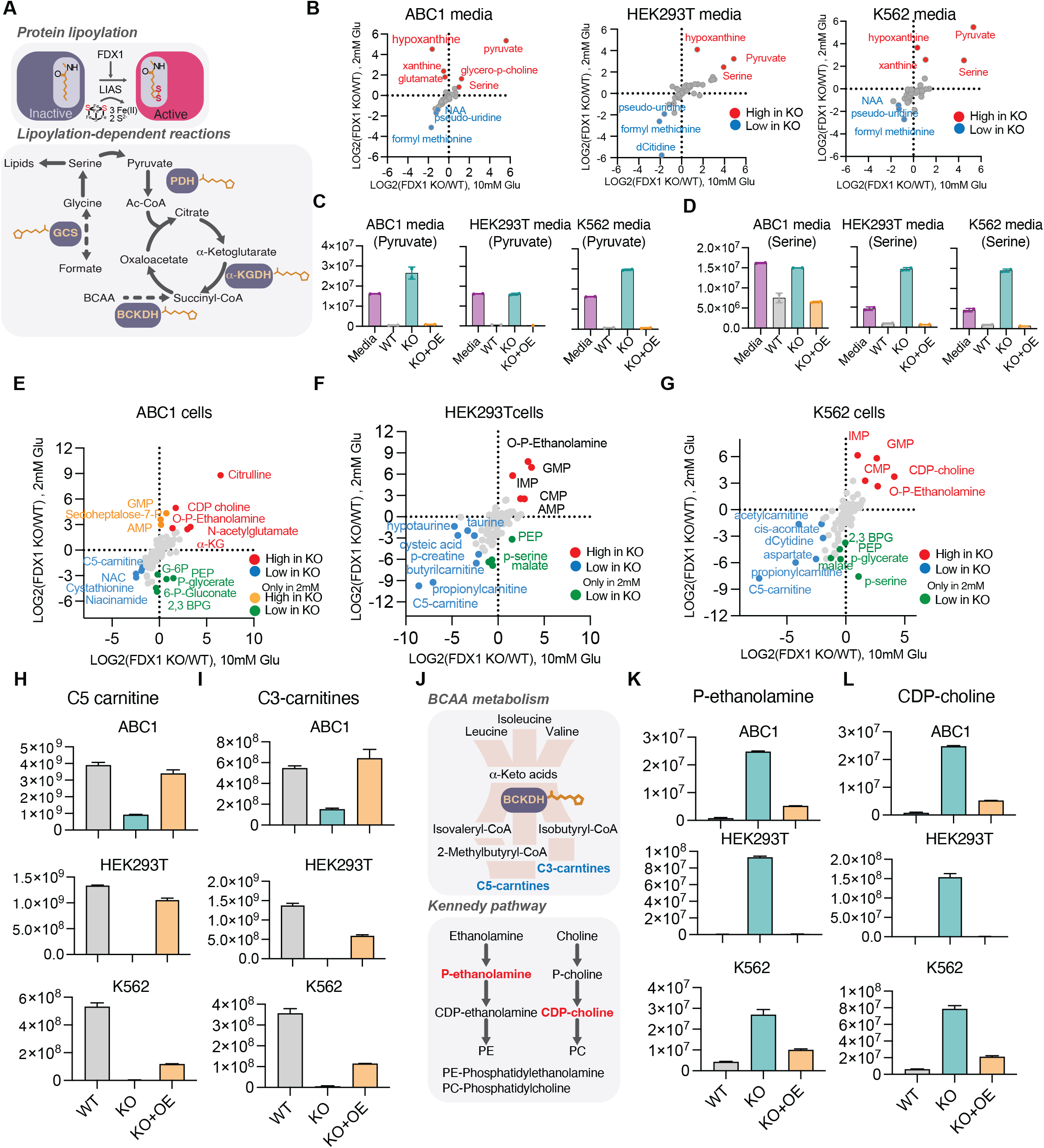
FDX1 loss-of-function results in metabolite signature consistent with protein lipoylation deficiency. (A) Schematic of the metabolic processes where lipoylated proteins are involved. (B) LOG2 fold change of metabolites in the media of either ABC1, HEK293T or K562 WT as compared to FDX1 loss-of function cells grown in media containing either 2mM (y-axis) or 10mM (x-axis) glucose. (C-D) changes in media pyruvate (C) or serine (D) levels in either WT, FDX1 loss-of-function (KO), or FDX1 loss-of-function cells with reconstituted FDX1 (KO+OE) HEK293T, ABC1 and K562 cells. (E-G) Intracellular metabolite level changes between WT and FDX1 loss-of-function cells (KO) ABC1 (E), HEK293T (F) and K562 (G) cells grown in media containing either 2mM (y-axis) or 10mM (x-axis) glucose. (H-L) Changes in intracellular levels of C5- (H) or C3- (I) carnitines or phospho-ethanolamine (K) or CDP-choline (L) in either WT, FDX1 loss-of-function (KO), or FDX1 loss-of-function cells with reconstituted FDX1 (KO+OE) HEK293T, ABC1 and K562 cells. Schematic of relevant pathways is presented (J).

The most significant intracellular metabolite change observed is the decrease in both C3- and C5-carnitine levels (Fig. 3H and I). C3- and C5-carnitines are downstream products of branched-chain amino acid (BCAA) metabolism that is mediated by the lipoylated BCKDH complex; the decreased levels in FDX1 KO cells suggest the BCKDH complex is inhibited in these cells (Fig. 3J). The a-ketoglutarate accumulation that is expected in the case of a-KGDH inhibition was only observed in ABC1 cells and not in HEK293T or K562 possibly due to alternative utilization of a-ketoglutarate in these cells. When cells were grown in low glucose conditions, these metabolic changes were further enhanced and a stronger decrease in C3- and C5-carnitines in FDX1-KO cells is observed (Fig. 3E-I), implying a failed attempt to channel branched-chain amino acid (BCAA) as an alternative fuel for the TCA cycle (37). Reduced levels of phosphoenolpyruvate (PEP) and other glycolytic intermediates such as 2,3-bisphosphoglyceric acid (2,3-BPG), and glucose-6-phosphate (G-6-P) were observed in FDX1-KO cells only when cultured in low glucose, likely reflecting altered metabolic flux.

An additional metabolite change that was observed in both low and normal glucose conditions was the dramatic increase in two intermediates (CDP-choline and phospho-ethanolamine) of the Kennedy pathway (Fig. 3K and L), that is responsible for the de novo synthesis of phosphatidylethanolamine and phosphatidylcholine (Fig. 3J). This FDX1-dependent increase in the Kennedy pathway intermediates suggests that there are global lipid metabolism alterations that might be induced due to the dysfunctional lipoylated TCA cycle complexes further explaining some of the observed dysregulation of lipid metabolism genes such as OCRL, LPCAT1 and SCD shown above (Fig. 2). Taken together, metabolite profiling of three cell lines following genetic perturbation of FDX1 and energy stress in the form of mild glucose deficiency, corroborate the hypothesis that FDX1 functions as part of the lipoylation pathway, and that its cellular role is manifested through the functions of the four lipoylation-dependent enzymes PDH, a-KGDH, BCKDH, and GCS.

### FDX1 binds LIAS in vitro and in cells

Given the essential role of FDX1 in regulating cellular protein lipoylation we next determined if FDX1 can directly bind LIAS. To test this, we implemented a Nano Luciferase (NanoLuc)-based complementation assay (38,39). In this assay two complementary fragments of NanoLuc (C1 and C2) are fused to two different proteins (A and B) of interest. The strength of the protein-protein interaction in cells can be quantified based on the intensity of luminescence (Fig. 4A) (38). One complementing fragment of Nanoluc was fused to the C-terminus of FDX1 (C2) and the other fragment (C1) was fused to other proteins of interest. To ensure this assay works for FDX1 as a mitochondrial protein, we show that as expected, FDX1 forms a strong dimerization interaction with itself but not with FDX2 that was used as a control (Fig. 4B). Using this complementation assay approach, we established that FDX1 can directly bind LIAS, GCSH and LIPT1 in cells but does not show significant binding to the lipoylated proteins such as DLAT, DLST or to other suggested FDX1 binding proteins such as ISCA2 and CYP11A1 (Fig. 4C). FDX2 was used as a control and showed no observed binding to the lipoylation regulating enzymes.

**Fig. 4.**
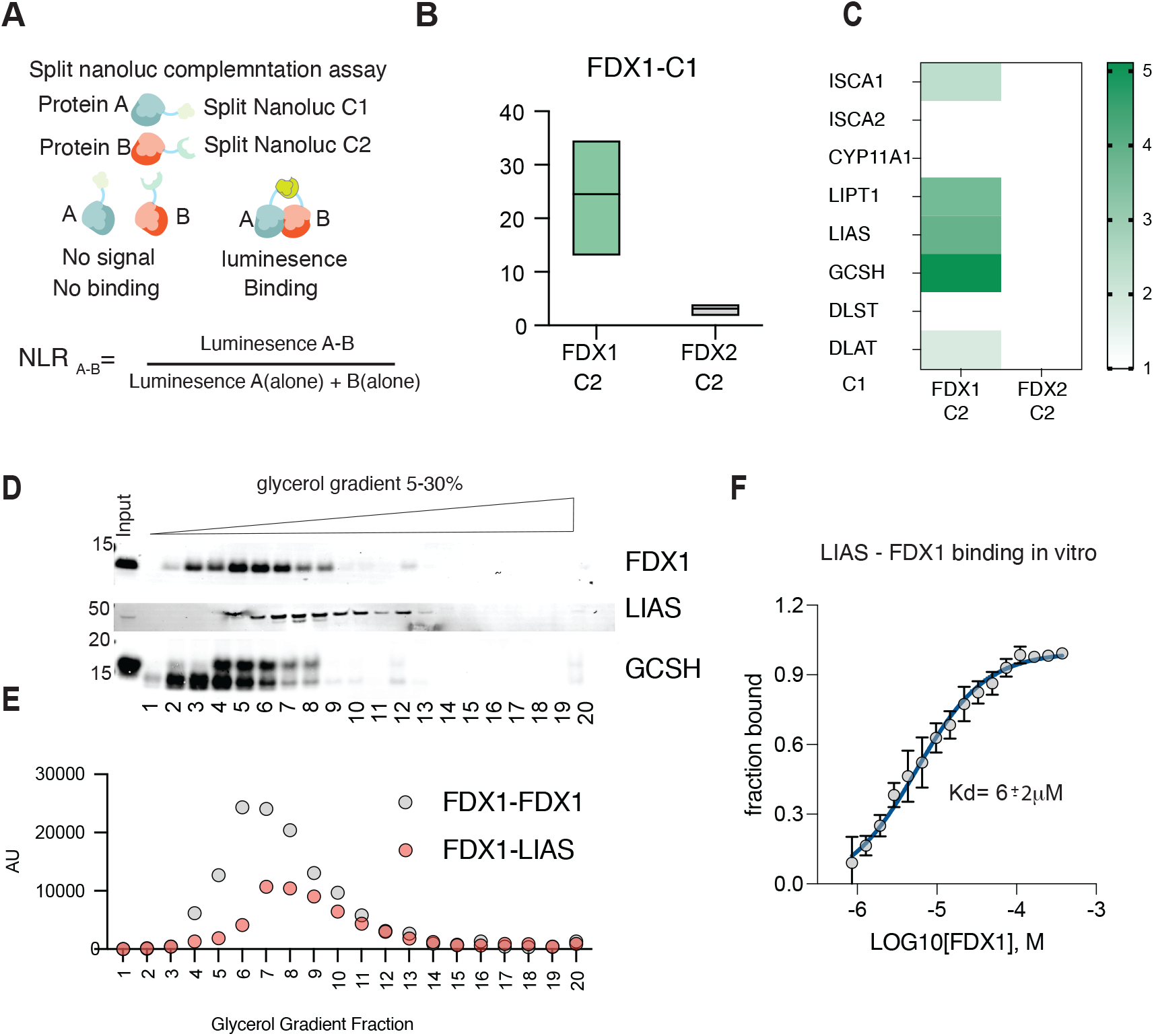
FDX1 directly binds LIAS in cells and in vitro. (A) Schematic describing the nano-luciferase complementation assays used and the equation used to normalize luminescence ratio (NLR) score corresponding to the raw luminescence value of the tested pair (A–B) divided by the sum of luminescence value from the two individual proteins (A-alone + B-alone). (B) The oligomerization of FDX1 was measured by overexpressing FDX1 tagged with fragment 1 of the complementation Nano-luciferase (C1) together with either FDX1 tagged with fragment 2 (C2) or FDX2 tagged with fragment 2 (C2) used as a negative control. Luminescence was measured 24 h after transfection (n = 3) (C) FDX1 or FDX2 used as a control tagged with fragment 1 of the complementation Nano-luciferase (C1) were overexpressed in the presence of indicated proteins tagged with fragment 2 of the complementation Nano-luciferase (C2). Luminescence was measured 24 h after transfection (n = 3). (B-C) Calculated NLR score is presented. (D) HEK293T cell lysates were subjected to a 5-30% glycerol gradient and FDX1, LIAS and GCSH protein sedimentation was analyzed using immunoblot analysis. (E) FDX1-C2 was overexpressed in HEK293T cells together with either FDX1-C1 or LIAS-C1 and lysates were analyzed for luminescence following sedimentation in 5-30% glycerol gradient. (F) MST results from titration of FDX1 to Red-NHS dye-labeled LIAS are shown, averaged raw data (n=3, black dots), the calculated binding curve fit from which the Kd was extracted (solid blue line), and the standard deviations (vertical black error bars). The fraction bound of labeled LIAS is plotted against the concentration of titrated FDX1 ligand and a K_d_ of 6 ± 2 μM was determined.

We were not able to show direct protein-protein interaction of FDX1-LIAS using an immunoprecipitation approach due to either the transient nature of the interaction or limitations of the antibody. This motivated us to confirm the direct binding of FDX1 to LIAS using two experimental approaches. First, using cellular extracts we show that endogenous FDX1, LIAS and GCSH co-migrate on a glycerol gradient (Fig. 4D) suggesting they might form a functional complex. Second, we performed the same glycerol gradient experiment on cells overexpressing either FDX1 or FDX1 with LIAS with complementary NanoLuc tags (C1 and C2) and determined that the peak of luminescence corresponds to the fractions where the endogenous proteins co-migrate (Fig. 4E). Third, we used a purified in-vitro experimental approach using a Microscale Thermophoresis assay (MST) that enables the characterization of binding without the need for immobilization of the proteins and can ensure the detection of weak or transient interactions (40). This MST binding approach revealed that purified FDX1 and LIAS enzymes (Fig. S3) bind in-vitro with a calculated Kd in the 6 μM range (Fig. 4F). Together these findings establish that FDX1 directly binds LIAS, supporting its enzymatic function of protein lipoylation.

## Discussion

In this work we explored the role and mechanism by which FDX1 regulates protein lipoylation in human cells. We were motivated by our recent finding that showed FDX1 is an upstream regulator of protein lipoylation in many cancer cell types (16). The goal of this study was first to establish if FDX1-regulation of protein lipoylation is direct, or indirect by inducing global Fe-S cluster biosynthesis defects. Second, to characterize the essentiality of FDX1-regulated lipoylation in cells growing in different glucose conditions. And third, to map the transcriptional and metabolic networks that are most affected by disruption of this poorly characterized pathway in human cells.

Our findings revealed that depletion of FDX1 results in a cell state with no detectable lipoylated proteins, minimal respiration and growth defects that are strongly affected by glucose availability in the media. FDX1 regulated protein lipoylation defects are not due to reduced Fe-S cluster biosynthesis. In fact, we saw no effect at all on the protein level of any of the tested Fe-S cluster proteins in cells with FDX1 loss-of-function. On the contrary, FDX1 loss-of-function resulted in the stabilization of LIAS protein as expected if FDX1 is an upstream regulator of LIAS activity. Inactive LIAS will not breakdown its auxiliary Fe-S cluster that is used as a substrate in the processes of lipoylation and as such will result in the stabilization of the enzyme. A recent report that was published during the preparation of this study showed in vitro that reduced FDX1 (by action of NADPH and FDXR) could efficiently replace dithionite as an electron source in the radical chain reaction of LIAS (13). These findings provide a possible mechanism of how FDX1 regulates protein lipoylation in cells. Our finding that FDX1 binds LIAS both in vitro and in cells suggests that the proximity of these enzymes and possibly of other lipoylation factors (GCSH, LIPT1) might be important for this reaction to occur efficiently in the mitochondria. Moreover, in this recent report (13) FDX1 regulated protein lipoylation was not associated with any observed changes in Fe-S cluster biosynthesis emphasizing that FDX1 is not involved in this process in human cells as previous findings suggested (8,9,11). Together, our initial observation that FDX1 regulates lipoylation in multiple cancers (16), the recently published mechanistic study of FDX1 regulation of lipoylation (13) and the work described above, provide compelling evidence on a new and important role of FDX1 in regulating protein lipoylation in human cells. Future work will determine if this is a newly acquired function of ferredoxins in evolution.

Our metabolomic and transcriptional data further suggest that regulating protein lipoylation is the predominant role of FDX1 in human cells. Metabolite profiling of cells with FDX1 loss-of-functions revealed reduced uptake of pyruvate and serine, reduced levels of BCAA breakdown products and in some cases accumulation of a-KG (ABC1 cells). All consistent with lipoylated protein complex function inhibition and reproducing previous metabolite assessment in models where lipoylated complexes were perturbed (36,41). Both the transcription and metabolite profiling experiments highlighted a dysregulation in lipid homeostasis for phosphatidylcholine (PC) in particular. In the FDX1 loss-of-function state there was an observed accumulation of the Kennedy pathway intermediates (CDP-choline and phosphoethanolamine) suggesting an impairment of the de novo synthesis of PC and an upregulation of LPCAT1 gene that can synthesize PC from LPC, an alternative pathway for PC synthesis. It remains to be determined if this dysregulation is due to a direct role of FDX1 in this process or due to global lipid metabolism rewiring as described in LIPT1 mutant cells (41). Moreover, our transcriptional data show the induction of both metabolism related genes and the integrated stress response (ISR) genes consistent with previous reports that indicated that the predominant gene signature activated by mitochondrial dysfunction is related to the ISR (33).

Together, our findings establish FDX1 as a key regulator of protein lipoylation through direct binding of LIAS, serving as a metabolic conditional lethality in cancer cells. It was surprising to reveal that despite the complete loss of protein lipoylation in FDX1 loss-of-function mutant cells, only a mild fitness defect was observed when cells were grown in high glucose conditions. Moreover, analysis of hundreds of cancer cell lines revealed that FDX1 loss-of-function is most correlated with LIAS loss-of-function and shows only modest cell growth inhibition effect in most cell lines tested (tested in normal media conditions). Only in a small subset of cancer cell lines the loss of FDX1 or LIAS is essential and results in complete growth inhibition (depmap.org). This is consistent with our transcriptional data where we observed the induction of the integrated stress response in normal growth conditions in K562 cells, whereas in ABC1 cells, the same stress response was observed only when cells were glucose deprived. These findings suggest that the dependency of cells on FDX1 and lipoylation pathways depends on both the intrinsic metabolic wiring of the cell and nutrient availability. Revealing the specific genetic and cellular states that might affect this dependency will establish if FDX1 regulated protein lipoylation is a unique vulnerability in a subset of cancers.

## Acknowledgments

We thank Joshua Dempster for help with the gene expression analysis. This work was supported by the National Cancer Institute grant 1 R35 CA242457-01 (TRG), by Riva therapeutics (PT, NRB and MBD), Research and Recruitment Funding by BCH (BP and NK), National Institutes of Health (awards GM-122595 to SJB.), the National Science Foundation (MCB-1716686 to SJB.), and the Eberly Family Distinguished Chair in Science (to SJB). SJB is an investigator of the Howard Hughes Medical Institute. NK is a Pew Biomedical Scholar.

## Author contributions

PT conceived this study. MBD, NRB and PT performed experiments and data analysis. BP and NK performed and analyzed the metabolomic experiments. DMW supervised by SJB performed the in vitro experiments. AC performed the gene expression analysis. PT and TRG supervised the study. PT wrote the manuscript with consultation from all authors.

## Conflict of interest

PT is a consultant and has received research funding from Riva Therapeutics.

## Materials and methods

### Correlation analysis

CRISPR *FDX1*, *FDX2* and *LIAS* dependency scores and *FDX1* and *CYP11A1* gene expression across 1128 cell lines from the Broad Institute’s public 22Q4 dependency map were correlated with other CRISPR dependency scores from the Broad Institute’s 22Q4 Dependency Map/CCLE release. Pearson’s correlations were performed in R using the cor.test function. P values were then corrected for false discovery using the Benjamini-Hochberg method of the p.adjust function in R and q-values were - log10-normalized. Top 1000 correlating genes were plotted using GraphPad Prism v8.3.0. Some of the results were used for gene enrichment analysis using STRING (42) version 11.5.

### Cell lines and culturing conditions

HEK293T, K562 and ABC1 cells used in this study were grown in RPMI media supplemented with 10% FBS and PS (for consistency between cell line experiments). For the glucose experiments cells were grown in RPMI without glucose (ThermoFisher Scientific, Waltham, MA) media that was supplemented with 10% dialyzed fetal bovine serum and 1mM Pyruvate and 2mM fresh glutamine and indicated concentrations of glucose (2mM-10mM). Uridine was not added to the media as it showed no effect on growth of any of the examined cell lines.

### Generation of FDX1 loss-of function cell lines

The general strategy for creating the FDX1 loss-of-function was to nucleofect a ribonucleoprotein (RNP) complex of CRISPR-Cas9 with the FDX1 targeting gRNA and then do single cell cloning to ensure maximal depletion of FDX1 gene expression. We could not grow our single cell clones of ABC1 cells so for these cells we undertook an additional step before the nucleofection step where ABC1 cells were pre-transduced with the previously validated gRNA targeting FDX1 (16) cloned into BRD016 guide only lentivector. Cells were selected for one week to ensure optimal gRNA expression and then subjected to the nucleofection protocol.

### Generation of FDX1 KO using nucleofection

The ribonucleoprotein (RNP) used in FDX1 KO by nucleofection was formed using a 1:1 ratio of Alt-R CRISPR-Cas9 tracrRNA (100 uM) and Alt-R CRISPR-Cas9 predesigned crRNA (100 uM) (Hs.Cas9.FDX1.1.AD IDT guide) (Integrated DNA Technologies, Newark, NJ). The mixture was incubated at 95°C for 5 minutes and then brought to room temperature to form the gRNA duplex. The gRNA duplex (50uM) was then combined with Alt-R Cas9 enzyme (61 uM) (Integrated DNA Technologies, Newark, NJ) at a ratio of 3:2 and incubated at room temperature for 10-20 minutes to form the RNP that were used for the nucleofection described below.

Nucleofection of ABC1, HEK 293T and K562 cells was conducted according to the manufacturer’s protocol (Lonza). 500,000 cells per sample were pelleted by centrifugation at 4°C using a tabletop centrifuge at 3000 rpm for 5 minutes. Cell culture media supernatant was aspirated and discarded, and the cells were resuspended in 30uL of a mixture containing 20uL Lonza SF cell line solution, 5uL FDX1 Cas9 RNP, 1uL electroporation enhancer and 4 uL PBS (Lonza Biosciences, Lexington, MA). Each cell suspension was added to individual wells of an electroporation cuvette (Lonza Biosciences, Lexington, MA), and then nucleofected using the Lonza 4D system program A549 pulse code CM-130. Cells were then transferred from the cuvette into pre-warmed media in a 6-well or 12-well culture plate and allowed to grow to 80% confluence. The HEK 293T and K562 nucleofected cells were further single cell plated to generate FDX1 KO single cell clones. ABC1 FDX1 KO cells could not be single cell cloned and the bulk nucleofected population was validated for high efficiency KO and used in subsequent experiments.

### Generation of the FDX1 reconstituted cell lines (KO+OE)

FDX1 was further introduced to the FDX1 loss-of-function cell lines clones selected to be used in the study. FDX1 was cloned into the pMT025 backbone vector (Addgene plasmid # 158579). Viral particles were produced in HEK293T cells and FDX1 loss-of-function cells (ABC1, HEK293T and K562) were infected and selected with puromycin to ensure homogenous FDX1 gene expression.

### Immunoblot analysis

Cells were lysed using RIPA lysis buffer (Sigma-Aldrich, St. Louis, MO) with protease inhibitor cocktail tablets (Sigma-Aldrich, St. Louis, MO) and then incubated on ice for 30 minutes to homogenize. Samples were subjected to centrifugation for 10 minutes at 10,000 RPM to pellet cell debris. Supernatant was collected for quantification and cell debris pellet was discarded. Protein was quantified using the bicinchoninic acid (BCA) method with a bovine-specific albumin standard curve for normalization. Each protein extract was boiled in 1x LDS sample buffer with 1:10 TCEP solution reducing agent and then size fractionated via pre-cast SDS-PAGE Bis-Tris 4-12% gels (Thermo Fisher Scientific, Waltham, MA), and transferred onto nitrocellulose membranes with the iBlot-2 system (Thermo Fisher Scientific, Waltham, MA). Each membrane was then incubated at room temperature for 1 hour in LICOR Odyssey blocking buffer then incubated at 4°C with the respective antibody 1 hour-overnight. The membranes were then washed 4 times with 1x TBS with 0.1% Tween20 (TBST) and incubated with fluorophore-specific IRDye secondary antibodies (LI-COR, Lincoln, NE). The membranes were then washed again 4 times with TBST before being imaged on a LI-COR Odyssey machine. Antibodies used in this study include: FDX1 (abcam ab108257), Lipoic acid (Millipore, cat# 437695 and abcam ab58724), LIAS (Proteintech 11577-1-AP), DLAT (Cell signaling 12362S and Thermo), DLST (cell signaling 5556S), Vinculin (abcam ab130007), CISD1 (Thermo Scientific 16006-1-AP), NTHL (Invitrogen PA316563), Actin (Cell signaling 8H10D10), ACO-2 (abcam ab129069), TOMM20 (abcam ab186735), POLD1 (abcam ab186407), GCSH (Proteintech 16726-1-AP)

### RNA extraction

RNA was extracted from cells using TRIzol™ (Thermo Fisher Scientific, Waltham, MA) according to manufacturer’s protocol.

### Library Preparation with PolyA selection and Illumina Sequencing

Library preparation and sequencing were conducted by Azenta life sciences according to the following protocol. RNA samples were quantified using Qubit 2.0 Fluorometer (ThermoFisher Scientific, Waltham, MA, USA) and RNA integrity was checked using TapeStation (Agilent Technologies, Palo Alto, CA, USA).

RNA sequencing libraries were prepared using the NEBNext Ultra II RNA Library Prep for Illumina using manufacturer’s instructions (New England Biolabs, Ipswich, MA, USA). Briefly, mRNAs were initially enriched with Oligod(T) beads. Enriched mRNAs were fragmented for 15 minutes at 94°C. First strand and second strand cDNA were subsequently synthesized. cDNA fragments were end repaired and adenylated at 3’ends, and universal adapters were ligated to cDNA fragments, followed by index addition and library enrichment by PCR with limited cycles. The sequencing libraries were validated on the Agilent TapeStation (Agilent Technologies, Palo Alto, CA, USA), and quantified by using Qubit 2.0 Fluorometer (ThermoFisher Scientific, Waltham, MA, USA) as well as by quantitative PCR (KAPA Biosystems, Wilmington, MA, USA).

The sequencing libraries were multiplexed and clustered onto a flowcell. After clustering, the flowcell was loaded on the Illumina instrument according to manufacturer’s instructions. The samples were sequenced using a 2×150bp Paired End (PE) configuration. Image analysis and base calling were conducted by the control software. Raw sequence data (.bcl files) generated from the sequencer were converted into fastq files and de-multiplexed using Illumina’s bcl2fastq 2.20 software. One mismatch was allowed for index sequence identification

After investigating the quality of the raw data, sequence reads were trimmed to remove possible adapter sequences and nucleotides with poor quality using Trimmomatic v.0.36. The trimmed reads were mapped to the reference genome available on ENSEMBL using the STAR aligner v.2.5.2b. BAM files were generated during this step. Unique gene hit counts were calculated by using feature Counts from the Subread package v.1.5.2. Only unique reads that fell within exon regions were counted.

The exploratory analysis used TPM-normalized counts of RNAseq data. The data was first filtered to only those genes who had counts greater than two (TPM > 2) in all conditions, and subsequently log-transformed. Each cell line was then analyzed individually.We centered and scaled the log(TPM+1)data such that each gene’s expression is centered at 0 with a standard deviation of 1. Using the cleaned and transformed data, we conducted Principal Component Analysis (shown in Figure 2), which revealed clear separations between samples.

After completing our exploratory analysis, we sought to identify which genes were significantly driving the separation seen in the PCA results. We began this analysis with the raw read counts and filtered to only those genes who had counts greater than zero in all conditions. Each cell line was again analyzed individually. Using the jackstraw method (43,44), we computed p-values indicating how significantly each gene contributed to the first two principal components. Using these results, we filtered to genes with p-values (PC weights) < 0.01. We then applied DESeq2 to this set of significant genes for each of our hypotheses. Our hypotheses were 1) H1-Given wild type, low glucose vs. high glucose, 2) H2-Given low glucose, FDX1 KO vs. wild type, 3) H3-Given high glucose, FDX1 KO vs. wild type, and 4) H4-Given FDX1 KO, low glucose vs. high glucose. Lastly, using the results from DESeq2, we conducted gene set enrichment analysis to determine the gene categories driving the PC weights under the specific experimental settings (H2 and H3).

### Metabolite Profiling by mass spectrometry for detection of polar metabolites

Cells from culture were washed with ice cold 0.9% NaCl briefly and extracted in buffer (80% Methanol, 25 mM Ammonium Acetate and 2.5 mM Na-Ascorbate prepared in LC-MS water, supplemented with isotopically labelled amino acid standards [Cambridge Isotope Laboratories, MSK-A2-1.2], aminopterin, and reduced glutathione standard [Cambridge Isotope Laboratories, CNLM-6245-10]) and transferred into 1.5ml tubes. Samples were vortexed for 10 sec, then centrifuged for 10 minutes at 18,000g to pellet cell debris. The supernatant was dried on ice using a liquid nitrogen dryer. Metabolites were reconstituted in 15 uL water supplemented with QReSS [Cambridge Isotope Laboratories, MSK-QRESS-KIT] and 1 uL was injected into a ZIC-pHILIC 150 x 2.1 mm (5 um particle size) column (EMD Millipore) operated on a Vanquish™ Flex UHPLC Systems (Thermo Fisher Scientific, San Jose, CA, USA). Chromatographic separation was achieved using the following conditions: buffer A was acetonitrile; buffer B was 20 mM ammonium carbonate, 0.1% ammonium hydroxide in water. Gradient conditions were: linear gradient from 20% to 80% B; 20–20.5 min: from 80% to 20% B; 20.5–28 min: hold at 20% B at 150 mL/min flow rate. The column oven and autosampler tray were held at 25 °C and 4 °C, respectively. MS data acquisition was performed using a QExactive benchtop orbitrap mass spectrometer equipped with an Ion Max source and a HESI II probe (Thermo Fisher Scientific, San Jose, CA, USA) and was performed in positive and negative ionization mode in a range of m/z = 70–1000, with the resolution set at 70,000, the AGC target at 1 × 10^6^, and the maximum injection time (Max IT) at 20 msec.

### Data Analysis for metabolomics

Relative quantification of polar metabolites was performed with TraceFinder 5.1 (Thermo Fisher Scientific, Waltham, MA, USA) using a 7 ppm mass tolerance and referencing an in-house library of chemical standards. Pooled samples and fractional dilutions were prepared as quality controls and injected at the beginning and end of each run. In addition, pooled samples were interspersed throughout the run to control for technical drift in signal quality as well as to serve to assess the coefficient of variability (CV) for each metabolite. Data from TraceFinder was further consolidated and normalized with an in-house R script: (https://github.com/FrozenGas/KanarekLabTraceFinderRScripts/blob/main/MS_data_script_v2.4_20221018.R). Briefly, this script performs normalization and quality control steps: 1) extracts and combines the peak areas from TraceFinder output.csvs; 2) calculates and normalizes to an averaged factor from all mean-centered chromatographic peak areas of isotopically labeled amino acids internal standards within each sample; 3) filters out low-quality metabolites based on user inputted cut-offs calculated from pool reinjections and pool dilutions; 4) calculates and normalizes for biological material amounts based on the total integrated peak area values of high-confidence metabolites. In this study, the linear correlation between the dilution factor and the peak area cut-offs are set to RSQ>0.95 and the coefficient of variation (CV) < 30%. Finally, data were Log transformed and Pareto scaled within the MetaboAnalyst-based statistical analysis platform (PMID: 25897128) to generate PCA, PLSDA, and heatmaps.

### Complementation assay

The vectors used for the Nano-Luc complementation assay was created by inserting the FDX1 and other genes of interest into N2H plasmids (Choi et al., 2019) (pDEST-C1 or -C2 kindly received from Yves Jacob, https://www.addgene.org/Marc_Vidal/) using the Gateway LR reaction. For the complementation assay 100ng of total DNA (1:1 ratio of plasmids) was transfected into 293T cells in 96-well plates using TansIT(R)-LT1 (Mirus) transfection reagent according to manufacturer’s protocol. Luminescence was measured 24-36 hours after transfection using Nano-glo (Promega) and viability (that was not affected) was measured using cell-titer glo (Promega) to ensure there were no viability differences (>10%) between the samples. All experiments were conducted with at least three biological replicates of each condition.

For the glycerol gradient, cells growing in 6-wells were transfected with FDX1-C2 and either FDX1-C1 or LIAS-C1 (2.5ug total plasmid at 1:1 ratio). 36 hours after transfection cells were lysed and subjected to glycerol gradient as described below.

### Glycerol gradient

The 5%-30% glycerol gradient was generated using the gradient master mixer (BioComp Instruments) with buffers containing 20 mM HEPES-KOH (pH 7.9), 150 mM NaCl, 1.5 mM MgCl_2_ and either 5% or 30% glycerol. HEK293T cells were lysed in lysis buffer (150 mM NaCl, 50 mM Tris-HCl, 1% NP-40, 1X Halt protease and phosphatase inhibitors). Cell lysates was spun at 20,000xg for 10 minutes at 4°C and supernatant was loaded onto the glycerol gradient. Gradients were spun at 45,000 RPM for 16 hours at 4°C in SW55Ti rotor using an ultracentrifuge (Beckman). After centrifugation, 250uL fractions were collected off the top of the glycerol gradient. For total protein lysates, NuPage sample buffer was added for a final concentration of 1X sample buffer and 16uL of sample was run on a 4-12% Bis-Tris gel following routine immunoblot procedure.

### Recombinant LIAS and FDX1 proteins over-production in *E. coli* and purification

Genes encoding LIAS (aa 28–372) and FDX1 (aa 61 – 184) lacking their respective mitochondrial transit peptides were custom synthesized at ThermoFisher Scientific after codon optimization for recombinant protein overexpression in *E. coli*. The LIAS gene was subcloned into a modified pSUMO vector (LifeSensors Inc.), (pDWSUMO), while the FDX1 gene was subcloned into pET28a vector. Both genes were cloned using *NdeI* and *XhoI* restriction sites of the appropriate vector. After gene sequence verification at Penn State Genomics Core Facility (University Park, PA), the plasmids were separately transformed into *E. coli* BL21 (DE3) cells carrying pDB1282, the plasmid which harbors an *isc* operon from *A. vinelandii* (45). As cloned, LIAS was over-expressed as a fusion with a ULP1 protease cleavable N-terminal SUMO tag that also carried a hexa-histidine tag at its N-terminus, while FDX1 was overproduced with an N-terminal hexahistidine tag. During purification, the SUMO tag was cleaved from the fusion protein affording pure LIAS with a Gly-His appendage at the N-terminus. Both proteins were overexpressed, purified and their respective iron-sulfur clusters reconstituted following procedures as previously published for LIAS with no further modifications (18).

### Microscale thermophoresis (MST)

To further confirm the interactions between FDX1 and LIAS in vitro, their binding affinity was measured via MST. For these experiments, a Monolith NT.115 Microscale Thermophoresis instrument (NanoTemper Technologies GmbH, München, Germany) was used. For the binding assays, Monolith RED-NHS 2^nd^ generation protein labeling kit (NanoTemper Technologies GmbH) was used to label LIAS and, all the labeling and binding steps of the experiment were performed in an anaerobic chamber (Coy Laboratory products, Grass Lake, Michigan). The labeling, binding and data analysis steps were carried out following the procedures as recommended by the manufacturer of the labeling kit. The MST experiments were performed in triplicate at room temperature in MST Buffer (PBS buffer (19mM Na_2_HPO_4_, 2mM KH_2_PO_4_, pH 7.4, 137mM NaCl, 2.5mM KCl) and with 300 nM final concentration of the labeled LIAS protein as the target, while FDX1 was serially diluted (375 – 0.009 μM) as the ligand. The LIAS-FDX1 mixtures were incubated for 30 min at room temperature after which were separately loaded into MST standard capillaries. The capillaries were then sealed and removed from the anaerobic chamber for binding measurements. Measurements were performed at 22 °C using medium MST power and 100% excitation power settings on the MO.control data collection software (NanoTemper Technologies GmbH). The data were analyzed and fit using the MO.Affinity Analysis software (NanoTemper Technologies GmbH) to determine the Kd of FDX1 binding to LIAS. The data was plotted using GraphPad Prism v8.3.0.

## Supplementary data

Table S1- Gene expression and dependency scores from CCLE.

Table S2- Gene expression of HEK293T and K562 cells (WT, FDX1 KO and FDX1 KO +OE)

Table S3- Metabolite analyses of HEK293T and K562 cells (WT, FDX1 KO and FDX1 KO +OE)

