## Supplemental figures S1-S3 for "FDX1 regulates cellular protein lipoylation through direct binding to LIAS"

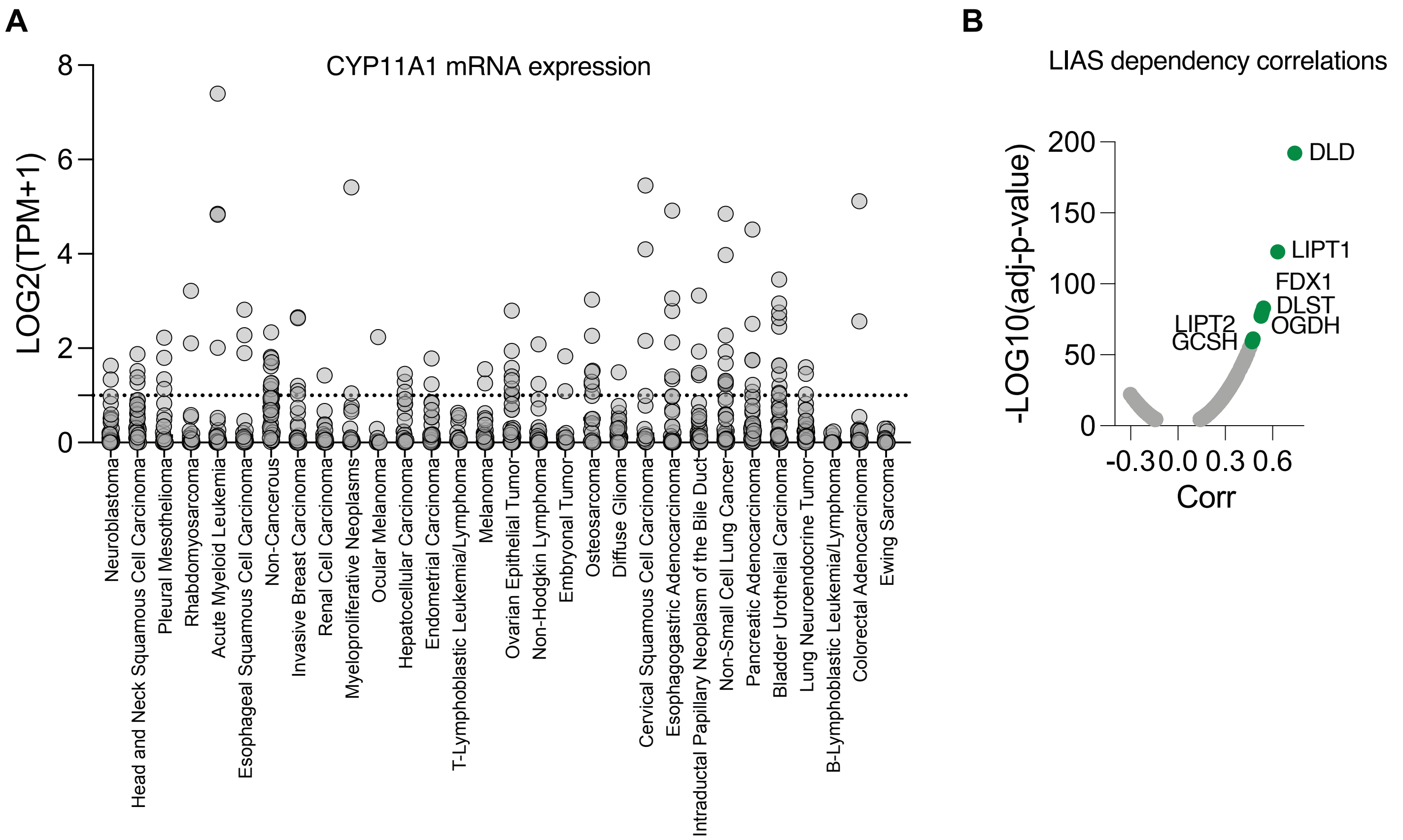


**Figure S1.** (A) CYP11A1 mRNA expression levels in cancer cell lines from the CCLE categorized by lineage. (B) The CRISPR/Cas9 LIAS loss-of-function viability outcome in 1128 cell lines correlated to the loss-of-function of all other tested genes. Plotted are the correlation values and the P-value for the 1000 most significant correlations. In green are genes associated with protein lipoylation pathway.

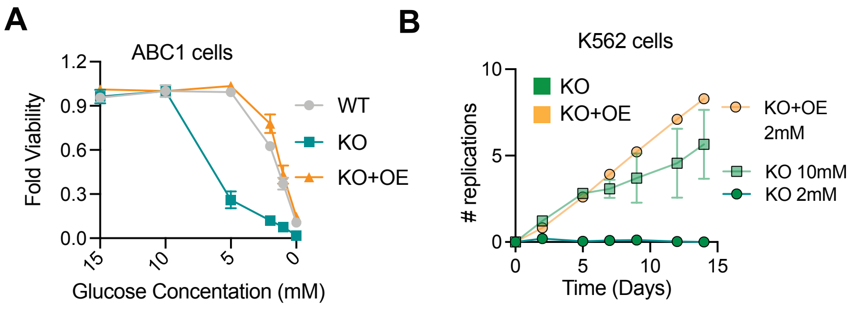


**Figure S2**. (A) Cell viability of ABC1 WT, FDX1 KO and FDX1 KO with FDX1 overexpression (KO+OE) 96 hours after growth in media containing either 2mM, 5mM or 10mM glucose. (B) prolifiration of FDX1 KO or FDX1 KO with FDX1 overexpression (KO+OE) cells was monitored for 14 days . Results represent the mean and standard deviation of at least three biological replicates.

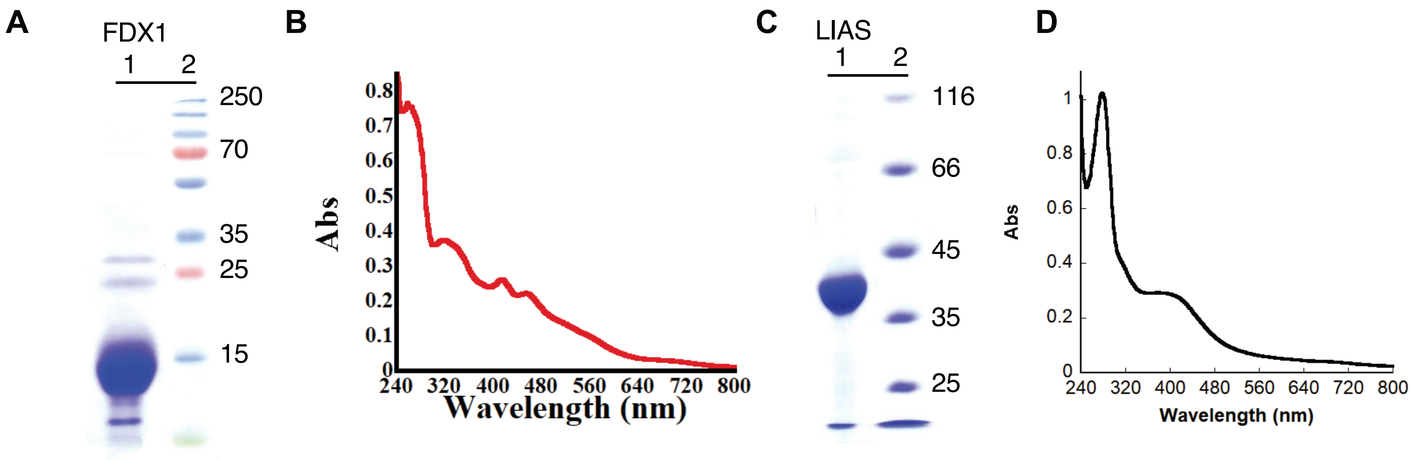


**Figure S3**. (A) SDS-PAGE gel analysis of the purified FDX1 (lane 1) and protein molecular weight markers (lane 2). (B) UV-vis absorption spectrum of 15 μM of the purified FDX1 protein. (C) pSDS-PAGE gel analysis of the purified LIAS protein (lane 1) and protein molecular weight markers (lane 2). (D) UV-vis absorption spectrum of 8 μM of the purified LIAS protein.
